# Nutritional analysis of commercially available, complete plant- and meat-based dry dog foods in the UK

**DOI:** 10.1101/2024.09.11.612409

**Authors:** R.A. Brociek, D. Li, R. Broughton, D.S. Gardner

**Affiliations:** School of Veterinary Medicine and Science, University of Nottingham, UK; School of Biosciences, University of Nottingham, UK; Institute of Aquaculture, University of Stirling, Scotland

**Keywords:** animal nutrition, canine, diet, dog, minerals, plant-based, vegan

## Abstract

**Background:** Adoption of a plant-based diet is a popular lifestyle choice for many owners of canine companion animals. Increasingly, owners would like to feed their canine companions a similar diet. A plant-based dietary pattern has been reported to be associated with some micronutrient deficiencies. Complete dog foods are, by definition, supposed to be nutritionally replete in all macro- and micronutrients. Few studies have reported a full nutritional analysis of complete, dry plant-versus meat-based dog foods.

**Method:** Here, 31 pet foods (n=19 meat-based, n=6 veterinary and n=6 plant-based) were analysed for total protein content and individual amino acids, fatty acids, major and trace elements, vitamin D and all B-vitamins.

**Results:** Nutritional composition of meat and plant-based foods were similar, except for iodine and B-vitamins, which were lower in plant-based foods. The majority (66%) of veterinary diets with lower total protein by design, were also deficient in one or more essential amino acids. Isolated instances of non-compliance to nutritional guidelines were observed across all food-groups. Of the tested nutrients 55%, 16%, 24% and 100% of foods met all amino acid, mineral, B-vitamin, and vitamin D guidelines, respectively.

**Conclusions:** Adopting a plant-based dietary pattern for your companion canine can provide nutritional adequacy with respect to the majority of macro- and micronutrients, but feeding supplemental iodine and B-vitamins should be considered. Veterinary diets, purposely low in crude protein, often have less than optimal essential amino acid composition. These data provide important new information for owners of companion canines being fed plant-based or veterinary diets.

## Introduction

Veganism is increasingly becoming a popular dietary choice for many people, whether it be for health reasons or concerns for animal welfare and/or the environment. The number of self-declared vegans in the UK quadrupled between 2014 and 2019, from 150,000 to 600,000 (0.25% to 1.2% of the population (<u>Society 2023</u>). Many share their homes with their omnivorous, canine companions. Owners of companion animals, who identify as vegetarian or vegan therefore face an ethical dilemma – should they feed animals to their animals? (<u>Bennett 2021</u>). Consequently, there has been an increase in the availability of ‘complete’ plant-based pet foods on supermarket shelves with little to no independent assessment of their nutritional soundness.

Meat-based food, including the incorporation of by-products from the meat industry, has long been seen as the ‘natural’ way to feed companion canines. Meat is high in protein and thus, provides the building blocks of protein via ‘proteogenic amino acids (AA)’, which are classified as either non-essential, conditionally-essential, or essential (EAA) (<u>Laidlaw and Kopple 1987</u>). The distinction being whether or not the body can form the amino acid from other substrates, for example by transamination (i.e. ‘non-essential’), whether the amino acid only becomes essential during certain high-demand ‘conditions’ such as pregnancy (‘conditionally-essential’) or whether the amino acids cannot be made in the body and thus must be acquired and ingested in the diet (‘essential’). In addition to protein, meat, dairy and other animal by-products tend to also be high in B-vitamins, selenium and organic phosphorous (<u>Pereira and Vicente 2013</u>). For such a diet to be labelled as ‘complete’ and to adhere to nutritional guidelines, in the UK/EU, administered by the Fédération Européenne de l’Industrie des Aliments Pour Animaux Familiers (FEDIAF – the European Pet Food Industry Federation) or, in the USA, by the Association of American Feed Control Officials (AAFCO), supplemental micronutrients are always added, as listed on the label. Dairy protein – casein – for example, is deficient in methionine, as are many other individual sources of plant-based protein; soybeans, beans and chickpeas are similarly unbalanced, lacking single amino acids from their profile, but when used in combination, can be effective for delivering sufficient amino acids and crude protein. Nevertheless, the few studies to date that have assessed the nutritional completeness of plant-based pet foods (sold in either Brazil (<u>Zafalon, Risolia, et al. 2020</u>) or Canada (<u>Dodd et al. 2021</u>)) reported multiple nutritional deficiencies in their composition.

In the UK, a survey of dog owners reported that the most important attributes any alternative diet would need to provide were ‘confidence about nutritional soundness’ and ‘confidence about pet health’ (cited by 84% and 83% of these respondents, respectively (<u>Knight et al. 2022</u>)). Similar observations were made in a separate study in North America; of those owners that did not already feed plant-based diets, a large proportion (45%; 269/599) were concerned, or wanted more information, about the nutritional completeness/adequacy of plant-based pet food (<u>Dodd et al. 2021</u>). Indeed, 74% stated that this was the primary reason that they didn’t currently feed a plant-based pet food. Other studies have drawn varying conclusions regarding the nutritional appropriateness of feeding companion canines a plant-based diet. One study concluded that further research was required to determine if long-term feeding of plant-based diets can meet and maintain amino acid and other nutrient targets in canines (<u>Cavanaugh et al. 2021</u>). Others found isolated instances of various micronutrient deficiencies in vegan foods available in Brazil (<u>Zafalon, Risolia, et al. 2020</u>). Further independent assessment is therefore warranted to provide evidence for owners of companion canines, that feeding a plant-based diet can provide nutritional completeness.

The primary objective of the current study was to measure the nutritional composition of complete, dry, meat- and plant-based foods available for canines on the UK market. We hypothesised that plant-based foods would be: 1) less ‘nutritionally-complete’ with lower compliance to nutritional guidelines than meat-based foods, 2) have lower protein and branched-chain amino acid content and 3) have lower B-vitamin, particularly B12, content. Foods indicated by a veterinarian to be fed to companion animals with kidney (e.g. chronic kidney disease, CKD) or urogenital problems (e.g. tendency to kidney stones) are, by design, low in protein but have a guaranteed analysis and are thus also labelled as ‘complete’. We hypothesised that 4) such ‘veterinary-renal’ foods would be low in essential amino acids relative to comparable ‘non-renal’ meat-based foods.

## Materials and Methods

### Selection of dog food

Thirty-one complete, dry dog food samples were acquired over a 4- week period (September 2022) from either online or high-street pet supermarkets in the UK, representing twenty-seven different brands. Exclusion criteria included not being a ‘complete’ food, not being labelled as for either ‘adult’ or ‘adult and senior’ dogs, foods that weren’t readily available to the public (i.e. required a prescription) and foods that did not come in packages of 3kg or less, to limit food waste. Foods were grouped according to their main protein source as listed on the label: “meat-based” (n=19; including, poultry (n=7), lamb (n=6) and beef (n = 6); “plant-based”, including vegan (n=4) and vegetarian foods (n=2). These six plant-based foods were the only brands available meeting our criteria sold in the UK, at the time of purchase. In addition, six guaranteed-analysis veterinary diets (n=6) were selected that were marketed as ‘reduced protein’, specifically for dogs with renal and/or urinary tract problems (‘Veterinary-renal’ foods). No specific flavour was declared for these foods.

### Preparation of dog food

Bags were inverted several times before opening. Approximately 100g of each dry food was sampled and frozen for a minimum of 24h at -20°C, before freeze-drying for 48-72h (Scanvac Coolsafe, Labogene, Denmark). Dry samples were then milled using a centrifugal mill (ZM 200 Ultra Centrifugal Mill & Vibratory Feeder DR100, Retzsch, Germany) at 10,000 RPM to a homogenous powder and stored in a 250ml polypropylene container at -20°C until required for analysis. Foods were not randomised or blinded for the authors. During preparation, all foods were given a sample number between 259 and 300 which was provided to analysts and technicians handling the samples, blinding these individuals to the provenance of each sample. All 31 complete dry dog foods were tested singularly and compared to European Pet Food Industry Federation (FEDIAF) guidelines for maintenance of adult dogs per 1000kcal of metabolisable energy (ME), at a maintenance energy requirement (MER) of 110kcal/kg^0.75^ body weight, necessary for dogs of moderate activity level (1-3hrs/day) (<u>FEDIAF 2022</u>).

### Protein and moisture content

Crude protein was determined using FlashEA® 1112 N/Protein (Thermo Fisher Scientific™) Nitrogen and Protein Analyzer, by directly measuring nitrogen content (<u>Krotz 2017</u>) and subsequently multiplying by 6.25. Directly-measured versus declared-on-label crude protein was then compared. Moisture content was determined by weighing a known quantity (100-200g) of frozen food, freeze-drying to completeness under vacuum for at least 2 days and re-weighing.

### Amino acid analysis, preparation of sample

Samples containing approximately 5mg nitrogen were oxidised in 20ml headspace glass crimp neck vials with 2.5ml of chilled, freshly made oxidation solution (10% of hydrogen peroxide (30% v/v) incubated (1 hour at 20-30 °C) in 87% v/v formic acid with 0.55% (w/v) phenol as an oxygen scavenger) for 16-18 hours at 4°C. After oxidation, 0.42g of sodium metabisulphite was added to decompose any excess oxidation reagent. The samples were then hydrolysed with 2.5ml 12M HCl and 0.5 ml of 6M HCL containing 1% phenol (w/v). The contents were mixed on a vortex mixer until all the sample was finely distributed in the acid. After mixing, the solutions were hydrolysed at 110°C for 24 h in an air draft oven. Sample pH was adjusted with 4M ammonium formate to 2.75 and made up to a volume of 50ml with 20mM pH2.75 ammonium formate buffer. After centrifugation at ×3000g for 2 minutes, 1-2 ml supernatant was passed through 0.22µm filter. The filtrate was diluted accordingly with ammonium formate buffer and an internal standard mix compromising cell-free stable isotope labelled (^13^C,^15^N) target amino acids to adjust the final amino acid values in the sample to be within the calibration range (nitrogen, 1-10µg/mL) of the instrument. A standard reference material of soy flour (National Institute of Standards and Technology, Maryland, USA, SRM 3234) was analysed in parallel to validate the accuracy of the hydrolysis and the amino acid analysis.

### Amino acid analysis; uHPLC-MS/MS

An aliquot of 200μL was dispensed into HPLC vials followed by 200μL of an internal standard comprising cell-free isotopically labelled (^13^C, ^15^C) target amino acids (Table S1). An amino acid standard curve was generated using Supelco amino acid standard mix, to which L-cysteic acid and methionine sulfone were added at a similar concentration with the amino acid standard mix. All samples were separated and analysed using a Thermo-Fisher Vanquish (uHPLC) and Altis Triple Quadrupole Mass spectrometer (MS/MS) with heated electrospray ionization (H-ESI) system. Positive ion mode was used for all amino acids. 1 µl was injected on a Thermo Scientific™ Acclaim™ Trinity P1 mixed mode column (150 mm x 2.1 mm, 3µM) at 30 °C. Mobile phases consisted of ammonium formate in water at pH 2.75 for phase A and a mixture of ammonium formate (100mM) in water and acetonitrile (80/20 v/v) for phase B. Chromatographic separation was achieved by gradient elution with MRM transition conditions as described (Table S2). Sheath gas was set at 45 arbitrary units, auxiliary gas at 15 arbitrary units, and spray voltage at 3500 V for positive ionization. Vaporizer temperature was set to 370 °C and transfer tube temperature to 270 °C, while source fragmentation was applied at 15 V. Data was acquired in Multiple Reaction Monitoring (MRM) mode using a resolution of 0.7 full width at half maximum (FWHM) for both quadrupoles. All compounds were detected in positive-ion mode.

### Amino Acid reporting

Nine out of ten essential amino acids were analysed in the current study, as tryptophan could not be detected using our current methods. Serine data are missing for nine samples, thus remaining data are reported for information only (supplementary Table S3). Additionally, the oxidation step of the preparation method leads to conversion of methionine to methionine sulfone and cysteine to cysteic acid ((<u>Manneberg, Lahm, and Fountoulakis 1995</u>), therefore results for methionine sulfone and cysteic acid will be reported as methionine and cysteine, respectfully. TraceFinder Version 4.1 was used to analyse raw data. Curve shape was standardised to ensure comparability between sample analyses. Conversion from reported units to g/1000kcal ME was completed using the calculated Atwater value for each individual food and the FEDIAF conversion value (<u>FEDIAF 2022</u>).

### Fatty acid methyl ester (FAME) lipid extraction and separation

5mL of 0.6M sucrose extraction buffer (for composition, see supplementary information, Table S4) was added to ∼2g dried food and homogenised using a GentleMACS tissue dissociator (Miltenyi Biotec, Ltd). 7mL of 0.6M sucrose extraction buffer was added to the homogenate, and the sample further centrifuged (Thermo Heraeus Pico 17, Thermo-Fisher Scientific™) at 3222rpm (2000*g*) for 5 mins. The homogenate was layered for sucrose cushion extraction (1mL 0.6M sucrose extraction buffer, 1.5mL homogenate, 2.5mL 0.25M sucrose extraction buffer) before ultracentrifugation at 100,000*g* for 1 hour (Hitachi CP80NX ultracentrifuge, P55ST2 rotor). The upper lipid layer was removed and further extracted by adding 5mL 2:1 chloroform:methanol, vortexed for 15 secs before the addition of 1mL 1% NaCl and further vortexed until homogenous. The sample was then centrifuged at 1000*g* for 2 minutes, and the lower (chloroform with lipid) fraction was transferred to a new glass centrifuge tube. Any remaining lipid was extracted by adding 3mL chloroform, vortexed for 20 seconds, centrifuged at 1000*g* for 2 minutes and the bottom fraction pooled to the new glass centrifuge tube. The pooled sample was dried under nitrogen and resuspended in 400µL of hexane. To this suspension, 0.7mL 10M KOH and 5.3mL methanol were added, and the sample heated at 55°C for 90min. After cooling, 0.58mL 12M H_2_S0_4_ was added and further incubated at 55°C for 90min. After cooling, 3mL hexane was added, mixed and centrifuged. The upper hexane layer was removed and concentrated by drying under nitrogen and reconstituting in 400µL hexane. The sample was stored at -30°C until required for analysis by GC-MS analysis.

### FAME Gas Chromatography Mass Spectrometry (GC-MS)

The fatty acid methyl esters (1µl) were injected (split ratio 50:1) into a gas chromatograph (GC) (Trace 1300, Thermo Fisher Scientific™) coupled with mass spectrometer (MS) (ISQ 7000, Thermo Fisher Scientific™). Separation of fatty acid methyl esters was performed with a Varian CP-Sil 88 (100m length, 0.25mm diameter, 0.20um film thickness, Agilent) capillary column with helium as carrier gas. Oven temperature (ramp up at 4°C/minute, from 140°C (hold for 5 minutes) to 240°C (hold for 10 minutes) and MS injector and transfer line temperature (260°C and 250°C, respectively) were preprogrammed. The ion source temperature set to 200°C. Characterization and identification of FAMEs was performed in scan mode. Quantification was completed by selective ion monitoring (SIM) mode of the most intense fragments. Data acquisition and processing were performed with the software Chromeleon (version 7.0, Thermo Fisher Scientific™).

### Mineral and trace elemental content analysis (ICP-MS)

Minerals and trace elements were determined by inductively coupled plasma-mass spectrometry (ICP-MS), expressed per unit dry weight (or Megacal), as previously described (<u>Alborough, Graham, and Gardner 2022</u>; <u>Davies et al. 2017</u>). Briefly, approximately 0.2 – 0.3g of sample and 0.1 – 0.2g bovine liver (as certified reference material, CRM: 1577C [National Institute of Standards and Technology (NIST)]) were digested using 3mL nitric acid, 3mL deionised (DI) water and 2mL H_2_0_2_ in a digestion microwave (Multiwavepro, Anton Parr, settings: 12 tubes, 1000W, 45 mins). Digesta was transferred into 50mL centrifuge tubes, and an additional 7mL deionised water, used to rinse any remaining sample. 500µL of digesta was pipetted into polypropylene ICP tubes before ICP-MS analysis using an iCAP-Q (Thermo Fisher Scientific™) by the Department of Environmental Science, Faculty of Science, Sutton Bonington Campus, University of Nottingham. Using this method, 32 major and trace elements are reliably reported, with n=13 referenced against FEDIAF guidelines (<u>FEDIAF 2022</u>). Standardisation between batches was achieved by adjustment to the CRM with n=23 elements reported.

### Vitamin analysis; Vitamin D

2-3g of freeze-dried food samples (n=29) were sent to The Institute of Aquaculture, University of Stirling, UK. Vitamin D was analysed by LC-MS/MS using a Waters Xevo TQ-S mass spectrometer coupled to a Waters Acquity I class UPLC with an Acquity UPLC BEH C18 column. Briefly, 600 mg of dry, homogenised pet food was weighed into glass vials, with 30 µl of a 2.5 µg/ml deuterated cholecalciferol (D3) and ergocalciferol (D2) standard added to each sample, plus calibration standards. A calibration curve was processed at the same time as the samples (0 to 50 µg/ml of non-labelled D2, D3). Briefly, vitamin D in the foods was extracted using 4 ml of 1.5 M potassium hydroxide in ethanol, with pyrogallol as the antioxidant for 1 hour at 80°C, followed by extraction with 3 ml of hexane, with the addition of 3 ml of 1% (w/v) potassium chloride solution. Hexane extracts were transferred to clean glass vials, dried under nitrogen, then re-constituted in ethyl-acetate. Samples were then derivatized using 4-phenyl-1,2,4-tirazoline-3,5-dione (PTAD) for 1 hour prior to analysis by LC-MS (see supplementary information, Table S5).

### Vitamin analysis; Vitamins B1 - 12

Approximately 2g of homogenised pet food (n=17) were sent to Creative Proteomics Ltd, USA, and analysed for the full range of B-vitamins using an AB Sciex QTRAP® 6500 LC-MS/MS platform. Briefly, each sample was further ground on a MM 400 mill mixer for 5 min at a shaking frequency of 30 Hz. 100 mg of the homogenised powder was weighed into a 5 mL tube and homogenized at 30 Hz for 5 min, followed by 5 min ultra-sonication in a water bath, then centrifuged at ×15,000*g* for 10 min. An internal standard (IS) of riboflavin (B2)-13C2/15N, nicotinamide (B3)-d4 and nicotinic acid (B3)-d4 was prepared in 65% acetonitrile. Serially diluted calibration solutions containing the 10 targeted vitamins were prepared in the IS solution. 10μL aliquots of the clear supernatants and the standard solutions were injected into a HILIC column (2.1*100 mm, 1.7µm) to run UPLC-MRM/MS with negative-ion mode on an Agilent 1290 UHPLC system coupled to an Agilent 6495C MS instrument, for detection and quantitation of ascorbic acid, nicotinic acid, vitamin B5 and B7, or with positive-ion mode for detection and quantitation of B9 and B12. For quantification of vitamin B1, B2 and nicotinamide, the sample solutions were diluted 10-fold with the IS solution before injection. The mobile phase was 2 mM ammonium acetate (A) and acetonitrile (B) for gradient elution (90% to 10% B in 12 min), at 0.3 mL/min and 40°C. Concentrations of the detected vitamins were calculated by interpolating the constructed linear regression curves of individual compounds, with the data acquired from injections of the sample solutions, in an appropriate concentration range for each metabolite (example trace for Vitamin B12, supplementary Figure S1). Limits of quantification for each of the B-vitamins are reported in supplementary Table S6.

### Statistical analysis

Data were analysed using analysis of variance (ANOVA) for the fixed effect of the three diet-types (meat-based or plant-based, veterinary). In order to meet assumptions for analysis by ANOVA, all data were checked for a normal distribution of residuals and respective Q-Q plots. If necessary, non-normally distributed data were log-transformed (log_10_) prior to analysis by ANOVA, or an alternative suitable non-parametric, distribution-independent test was used (e.g. Kruskall-Wallis NP-ANOVA). All such data were analysed using GraphPad Prism v9.5.0 (GraphPad Software Inc., California, USA) and GenStat v22 (VSNi Ltd., Rothamsted, UK). Since multiple amino acids, fatty acids and trace element data were derived from each sample, many may often show concordance between analytes in the same sample. To mitigate such over-dispersion, multivariate, linear discriminant analysis was used as an objective means to effectively demonstrate significant patterns in complex (i.e. multiple variates), potentially non-independent data using orthogonal partial least squares-discriminant analysis (OPLS-DA; SIMCA-P v19, Umetrics, Umea, Sweden). Certain analyses and graphical representations were also conducted in the open-source software JASP Team (v0.17.1.2023; jasp-stats.org), as indicated in appropriate Figure or Table legends.

### Data availability

All anonymized data for the products used in this manuscript are available from The University of Nottingham research data repository at http://doi.org/.

## Results

### Protein and amino acid content of foods

Direct analysis of crude protein content of all foods indicated similarity in protein content between meat- and plant-based foods, with veterinary foods being, by design, lower in total protein (Figure 1a). The directly measured, versus stated protein content on the label, corresponded well (Figure 1b). Measurement of all individual amino acids, including 9/10 essential for canines (Arginine, Histidine, Isoleucine, Leucine, Lysine, Methionine, Phenylalanine, Threonine and Valine) are also reported and data, as expected, were similar to total protein (Figure 1c). Compared across food types, then veterinary-renal diets generally had significantly reduced content of all amino acids (Table 1, for values per 100g DM). Nevertheless, essential amino acids are essential and must meet minimum inclusion levels; it was therefore notable that while 17/31 (55%, n=11/19 meat-based, n=2/6 veterinary, n=4/6 plant-based) foods met EAA minimum inclusion levels, many veterinary-renal foods did not (Figure 1e) – four of six being below nutritional guidelines for essential amino acids, with threonine below guideline inclusion levels in all four (range: 1.04-1.14g/1000kcal, FEDIAF: 1.3g/1000kcal; Figure 1e). Remarkably, one of the six veterinary-renal foods was low in 6/8 essential amino acids (isoleucine, leucine, methionine sulfone, phenylalanine, threonine and valine; Figure 1e). In an unbiased, multivariate analysis of all foods and all measurable amino acids, the three food types were clearly distinguishable from each other (Figure 1d). Most of the variation (68%) was explained by less total protein and individual amino acid content in veterinary foods (Figure 1e).

**Figure 1.**
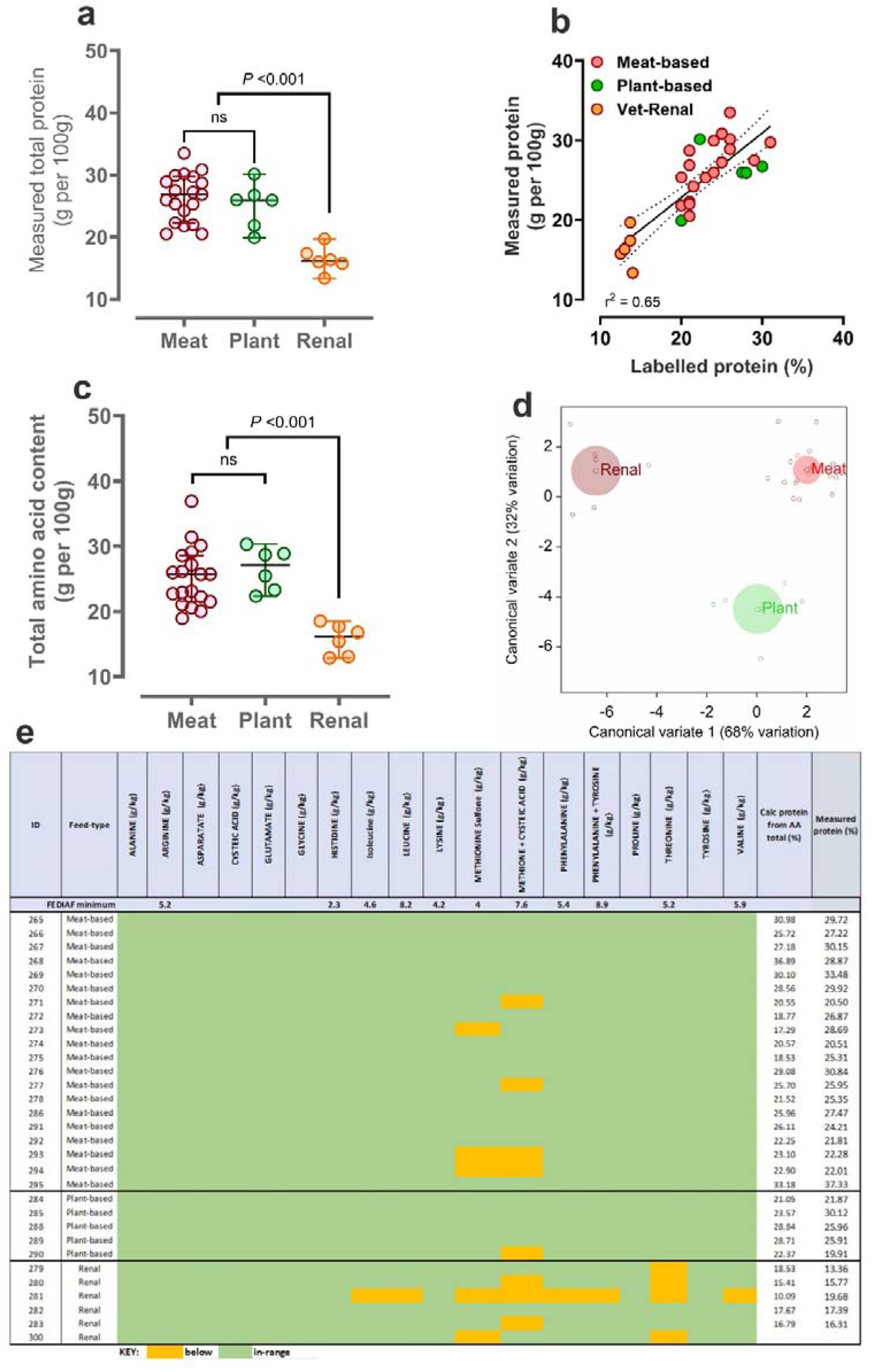
Protein and amino acid content of dry feeds for dogs according to feed-type. **a)** Individual data points for directly measured total protein, **b)** relationship between measured and labelled total protein, **c)** directly measured total alpha amino acids according to the three feed types, **d)** multivariate analysis (discriminant plot) with all feeds (n=31) and all measured amino acids (n=19) represented, **e)** data expressed relative to the guideline nutritional minimum for each individual amino acid (‘FEDIAF minimum’. **Green** boxes represent values in range according to our analyses, **yellow** boxes are below nutritional minimum.

**Table 1.** Amino acid content of dry feeds for dogs according to feed-type, per unit dry matter.

All data are mean ± SD, units per 100g DM, based on 110kcal/kg maintenance energy requirement (MER).
| **Amino acid** | **FEDIAF**<br>(per 100g DM) | **Meat-based**<br>(n=19) | **Plant-based**<br>(n=6) | **Renal**<br>(n=6) | **Statistics**<br>*P-value |
| --- | --- | --- | --- | --- | --- |
| Alanine (g) | - | 1.47 ± 0.06<sup>a</sup> | 1.31 ± 0.11<sup>a</sup> | 0.77 ± 0.11<sup>b</sup> | <0.001 |
| Arginine (g) | 0.52 | 1.50 ± 0.06<sup>a</sup> | 1.46 ± 0.11<sup>a</sup> | 0.73 ± 0.11<sup>b</sup> | <0.001 |
| Aspartate (g) | - | 2.28 ± 0.12<sup>a</sup> | 2.51 ± 0.22<sup>a</sup> | 1.44 ± 0.22<sup>b</sup> | 0.02 |
| Cysteic acid (g) | - | 2.93 ± 0.75<sup>a</sup> | 3.62 ± 0.42<sup>b</sup> | 2.53 ± 0.36<sup>a</sup> | 0.004 |
| Glutamate (g) | - | 3.58 ± 0.15<sup>a</sup> | 4.41 ± 0.27<sup>a</sup> | 2.16 ± 0.27<sup>b</sup> | <0.001 |
| Glycine (g) | - | 2.22 ± 0.03<sup>a</sup> | 1.44 ± 0.18<sup>a</sup> | 0.95 ± 0.18<sup>b</sup> | <0.001 |
| Histidine (g) | 0.23 | 0.49 ± 0.02<sup>a</sup> | 0.58 ± 0.03<sup>a</sup> | 0.32 ± 0.03<sup>b</sup> | <0.001 |
| Isoleucine (g) | 0.46 | 0.89 ± 0.04<sup>a</sup> | 0.99 ± 0.07<sup>a</sup> | 0.59 ± 0.07<sup>b</sup> | <0.001 |
| Leucine (g) | 0.82 | 1.85 ± 0.11<sup>a</sup> | 2.10 ± 0.20<sup>a</sup> | 1.32 ± 0.20<sup>b</sup> | 0.03 |
| Lysine (g) | 0.42 | 1.28 ± 0.07 | 1.32 ± 0.12 | 0.99 ± 0.12 | 0.12 |
| Methionine sulfone (g)<sup>1</sup> | 0.40 | 0.49 ± 0.02 | 0.49 ± 0.04 | 0.45 ± 0.04 | 0.77 |
| Methionine + Cysteic acid (g)<sup>1</sup> | 0.76 | 0.78 ± 0.03 | 0.85 ± 0.06 | 0.71 ± 0.06 | 0.23 |
| Phenylalanine (g) | 0.54 | 1.01 ± 0.05<sup>a</sup> | 1.19 ± 0.09<sup>a</sup> | 0.62 ± 0.08<sup>b</sup> | <0.001 |
| Phenylalanine + Tyrosine (g) | 0.89 | 1.64 ± 0.08<sup>a</sup> | 2.05 ± 0.14<sup>a</sup> | 1.03 ± 0.14<sup>b</sup> | <0.001 |
| Proline (g) | - | 1.67 ± 0.07<sup>a</sup> | 1.42 ± 0.14<sup>a</sup> | 0.74 ± 0.14<sup>b</sup> | <0.001 |
| Threonine (g) | 0.52 | 0.84 ± 0.04<sup>a</sup> | 0.87 ± 0.07<sup>a</sup> | 0.46 ± 0.07<sup>b</sup> | <0.001 |
| Tyrosine (g) | - | 0.63 ± 0.03<sup>a</sup> | 0.85 ± 0.06<sup>a</sup> | 0.42 ± 0.06<sup>b</sup> | <0.001 |
| Valine (g) | 0.59 | 1.20 ± 0.05<sup>a</sup> | 1.23 ± 0.08<sup>a</sup> | 0.69 ± 0.08<sup>b</sup> | <0.001 |

### Fatty acid content of foods

Broadly, the fatty acid composition of meat-based and veterinary foods was similar. Meat-based foods had the highest proportion of animal-based saturated fatty acids such as palmitic, stearic and arachidic acid (sum of saturated fats; meat-based, 67.2 ± 19.7; plant-based, 42.2 ± 5.1; veterinary, 56.6 ± 11.5 gms %; *P*=0.01). Plant-based foods had the highest incorporation of mono- and poly-unsaturated fatty acids; oleic, linoleic and linolenic acid (23.9 ± 12.5, 27.4 ± 8.6 and 3.43 ± 3.44 gms %, respectively).The majority of individual fatty acids in the pet foods analysed in this study were either saturated; C8:0 (caprylic – 23% of total), C16:0 (palmitic – 22% of total), C18:0 (stearic – 13% of total), C20:0 (arachidic - 1% of total) or unsaturated; C18:1n9 (oleic – 16% of total), C18:2n6c (linoleic – 14% of total), linolenic (C18:3n3 - 2% of total; Figure 2a). Whilst measurable, the combined sum of C6:0 (caproic), C10:0 (capric), C11:0 (undecanoic), C12:0 (lauric), C13:0 (tridecanoic), C14:1 (myristoleic), C15:0 (pentadecanoic), C17:0 (heptadecanoic), C18:1n9t (elaidic), plus other long-chain fatty acid derivatives (C20:0, arachidic – C24:1, nervonic) comprised <5% of total fat in each sample (‘other’ in Figure 2a).

**Figure 2.**
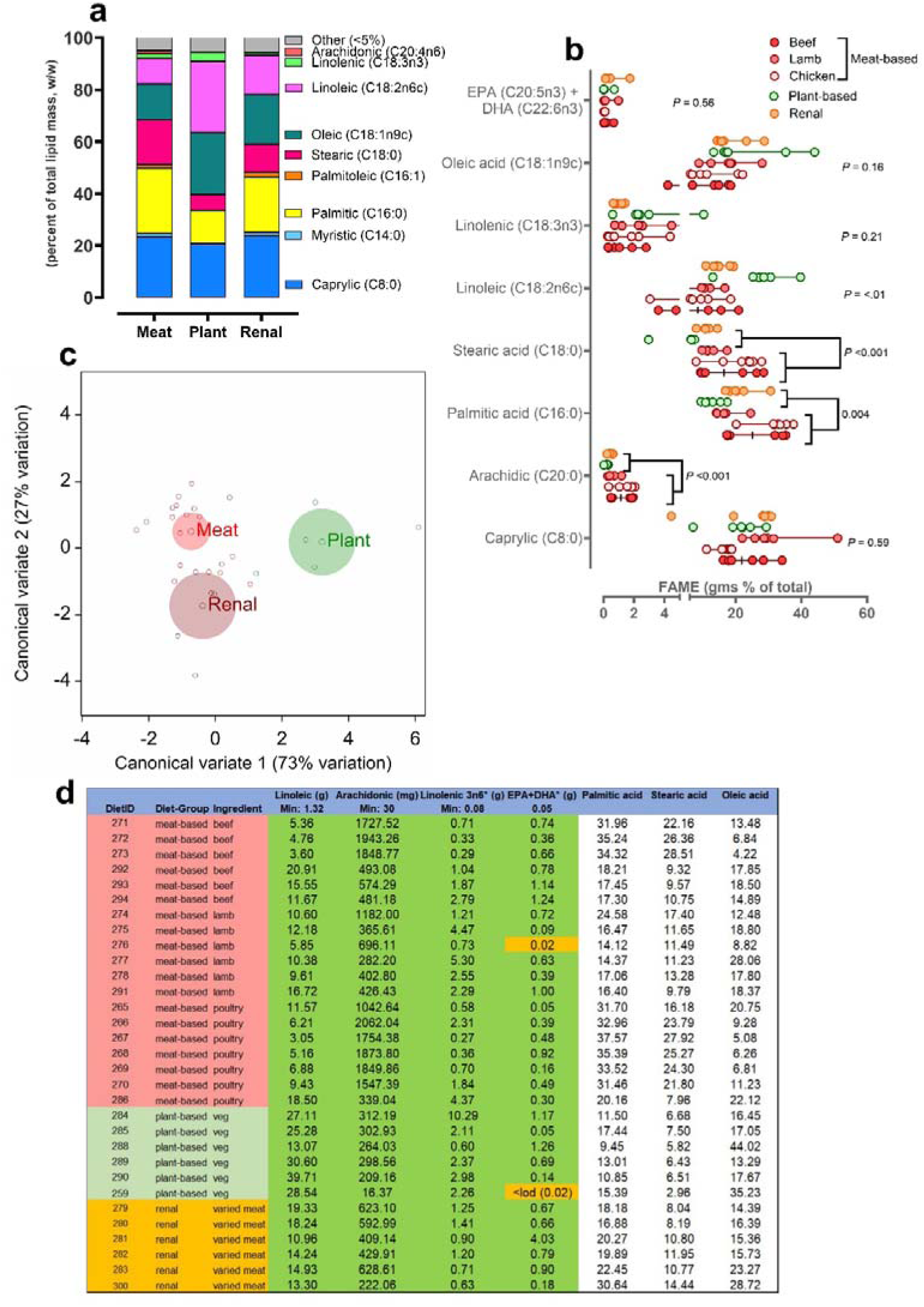
Fatty acid content of dry feeds for dogs according to feed-type. **a)** relative proportions (w/w %) of prevalent fatty acids between feed types, **b)** individual data points for measured fatty acids between feed types with beef, lamb and chicken combined as ‘meat-based’ for analysis by 1-way ANOVA, **c**) multivariate analysis (discriminant plot) with all feeds (n=31) and all measured fatty acids ≥LOD (n=9) represented, **d)** data (individual measured value) expressed relative to the guideline nutritional minimum for each individual fatty acid (‘Min’). **Green** boxes represent values in range according to our analyses, **yellow** boxes are below nutritional minimum. White boxes, no specified nutritional range

All foods were replete in linoleic acid, according to the nutritional guidelines (FEDIAF), on a gram per 100g total lipid (i.e. g % lipid mass) or mass-basis (i.e. ≥1.32g/100g DM; Figure 2b,d). Plant-based foods had significantly greater (*P* <0.01 by Kruskall-Wallis NP ANOVA) linoleic acid (27.3 ± 8.6 gms %) than meat-based (9.89 ± 5.17 gms %) and veterinary (15.2 ± 3.1 gms %) foods (Figure 2b). Unbiased multivariate discriminant analysis showed a clear separation of plant-based from both meat-based and veterinary – which were similar with respect to fatty-acid composition – along the first principle component (73% variation explained) distinguishing plant-based diets as having an overall greater incorporation of caprylic (+1.24 contribution to latent vector 1) and linolenic acid (+0.26; Figure 2c). Using macronutrient data on the labels of each food to calculate gross and metabolisable energy, according to Atwater criteria, indicated that veterinary foods had higher energy content than both meat or plant-based foods (Energy density: meat-based, 332 ± 16; plant-based, 328 ± 5; veterinary, 376 ± 10 kcal ME /100 g DM; *P*<0.001), largely due to incorporation of more fat in the food (% fat on label: meat-based, 12.8 ± 3.0; plant-based, 10.4 ± 1.8; veterinary, 17.3 ± 1.7 gms fat as fed, *P*<0.001).

### Major and trace minerals

Whilst only 16% of individual foods tested (n=5/31, all meat-based) satisfied all mineral guidelines, overall compliance was moderate-to-high (n=342/391 [87%] of minerals were ‘in-range’, i.e. n=13 guidelines × n=31 foods = 391 in total, excluding Ca and P which are purposely reduced in renal foods; Figure 3a). The individual foods with ‘out-of-range’ minerals tended to be isolated instances (e.g. of chloride or zinc), but below recommended incorporation of iodine and selenium were common in all foods (Figure 3a, Table 2), whether expressed per unit mass (Figure 3b) or unit energy (Figure 3c,d). Veterinary diets formulated for dogs requiring renal support were lower in calcium (*P*=0.009), phosphorous (*P*=<0.001), magnesium (*P*<0.001), iron (*P*=0.006) and selenium (*P*=0.01; Table 2). Plant-based foods had greater potassium and lower iodine content than other food types (both P ≤0.01).

**Figure 3.**
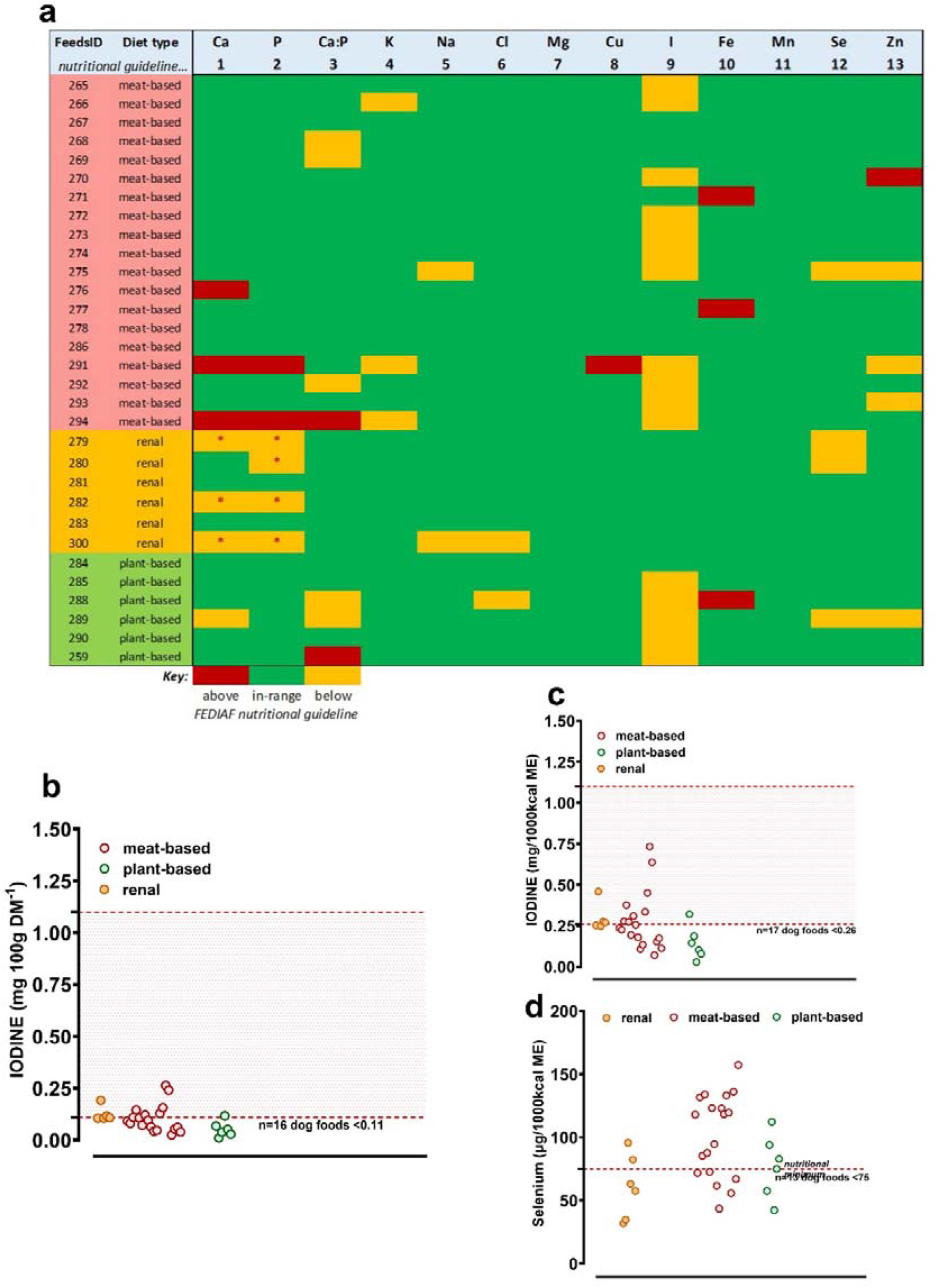
Major and trace elemental content of dry feeds for dogs according to feed-type. **a)** data expressed relative to the guideline nutritional minimum for each individual major and trace element (‘Nutritional Guideline’; respective FEDIAF guideline #1 - #13). **Green** boxes represent values in range according to our analyses, **yellow** boxes are below nutritional minimum, **red** boxes are above nutritional (or Legal) maximum. **yellow** boxes with a **\*** are allowable low values according to EU2020/354 intended use of feed for particular nutritional purpose (PARNUT). **b,c,d**, individual data points for measured iodine (**b,** mg/100g DM or **c**) mgs/1000kcal) and **d**) selenium between feed types with beef, lamb and chicken combined as ‘meat-based’. Upper and lower dashed lines give relevant nutritional range for compliance.

**Table 2.**
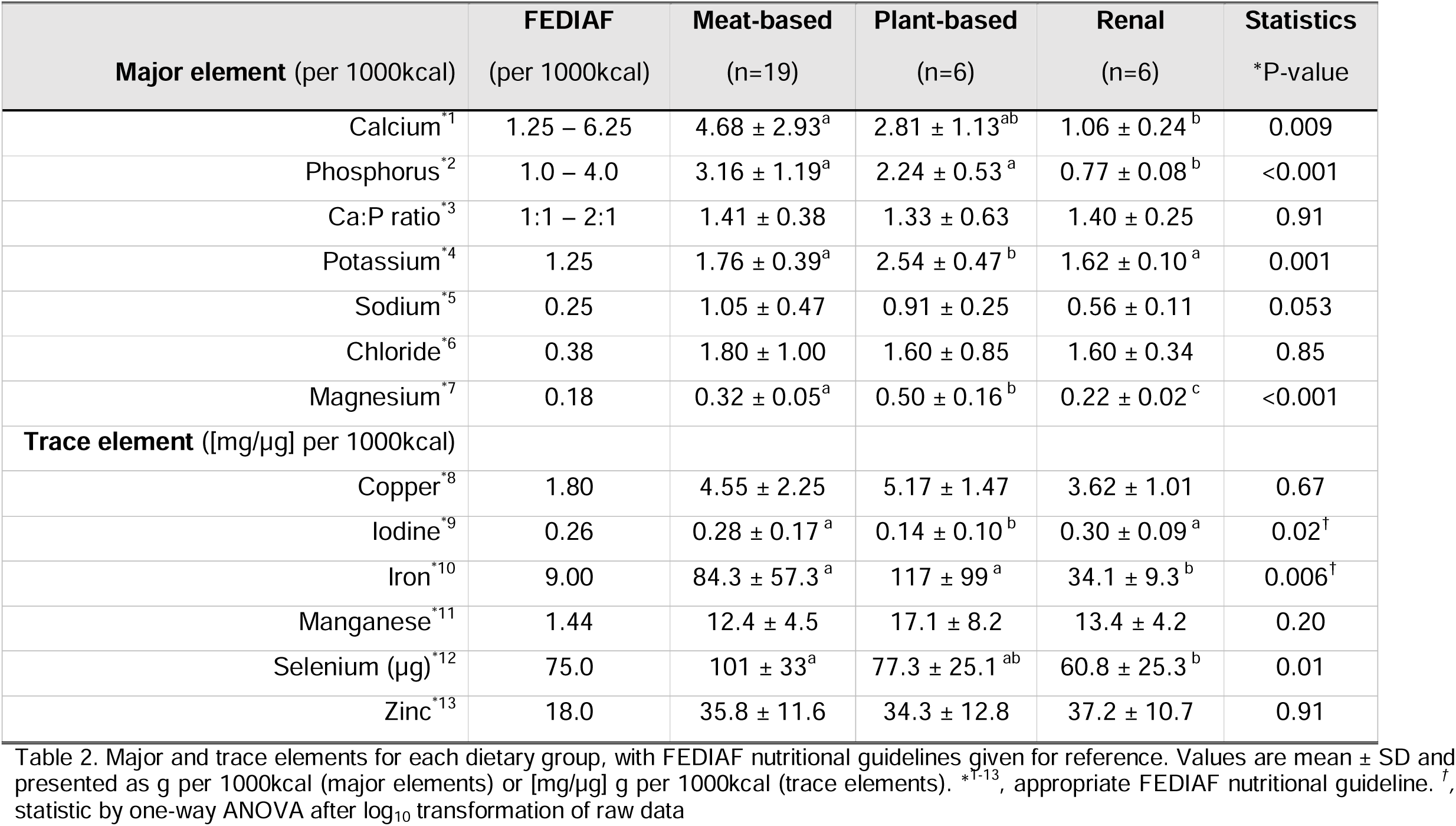
Major and trace elemental content of dry feeds for dogs according to feed-type.

### Vitamins, Vitamin D

All foods, when directly measured for vitamin D content, were within the recommended nutritional range (i.e., between FEDIAF nutritional minimum and maximum of 138-800 IU/1000kcal; Figure 4a). *B-vitamins*: 17 foods (meat-based, n=8; plant-based, n=5; veterinary, n=4) were analysed for a full panel of B-vitamins (B1, B2, B3, B5, B6, B7, B9 and B12), of which seven (all except vitamin B7) have FEDIAF nutritional minimum recommendations. B-vitamin content of foods were mostly comparable between food types, but consistently lower B-vitamin content was noted in plant- versus meat-based foods for vitamins B3, B9 and B12 (Table 3; Figures 4b,c; all at *P* ≤0.05, 1-way NP ANOVA). Accordingly, when summated, plant-based foods had lower B-vitamin content than meat- based foods (Figure 4d, Table 3). Overall, compliance of foods to nutritional recommendations for B-vitamins was poor – only four of 17 tested (23%) met all minimum requirements for B-vitamins, with any deviation from recommended being below the guideline level (Figure 4e).

**Figure 4.**
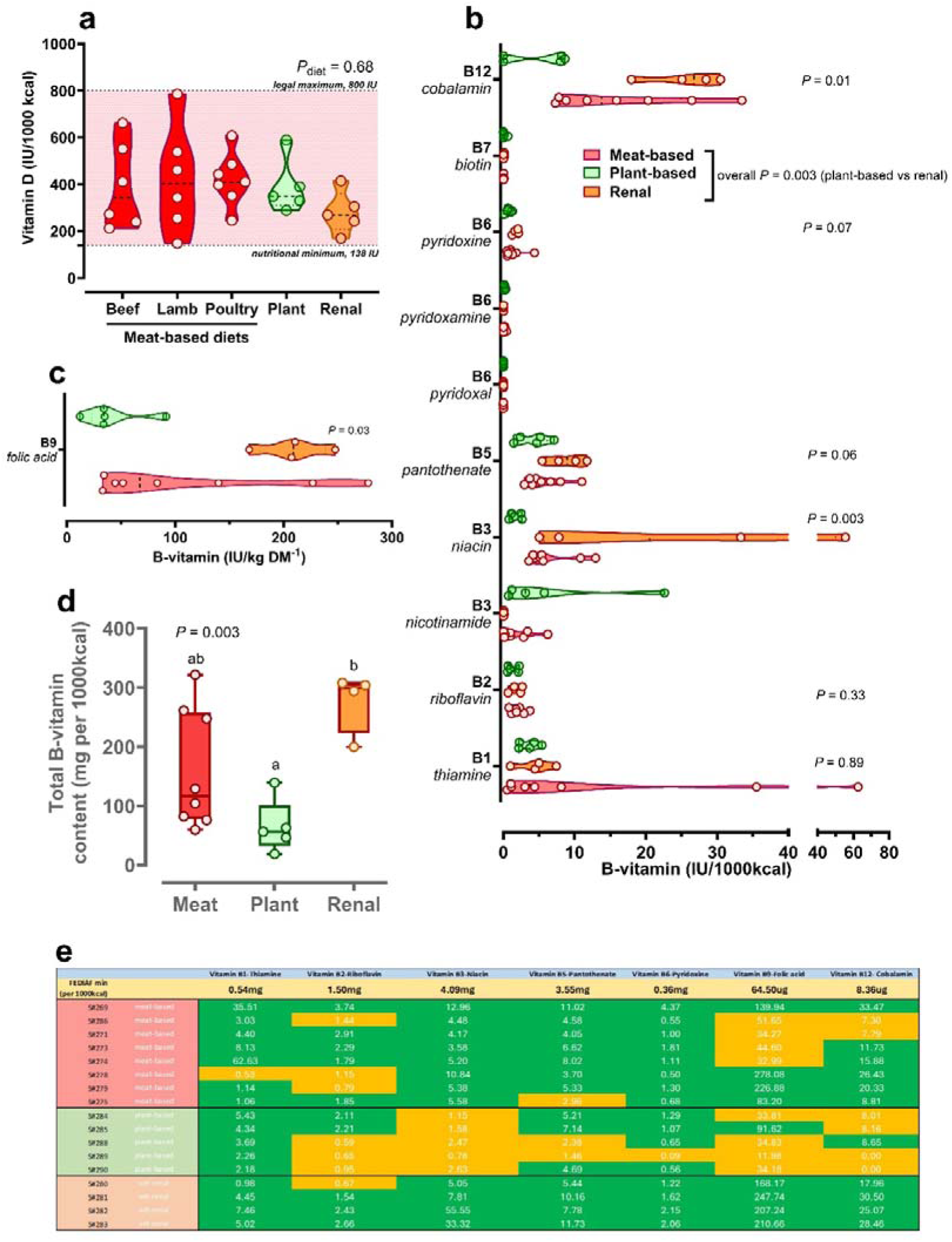
Vitamin content (B1-B12, D) of dry feeds for dogs according to feed-type. **(a)** individual data points for measured vitamin D between feed-types, **(b)** individual data points for measured b-vitamins in food types, *P*-value by NP ANOVA (Kruskall-Wallis test), **(c)** individual data points for measured vitamin B9 (folic acid) in food types, note different scale on x-axis, **(d)**, summated total B-vitamins between foods, *P*-value by NP ANOVA (Kruskall-Wallis test), **(e)**, data expressed relative to the guideline nutritional minimum for each individual B-vitamin (top row). **Green** boxes represent values in range according to our analyses, **yellow** boxes are below nutritional minimum.

**Table 3.** B-vitamin content of dry feeds for dogs according to feed-type.

All data are expressed as units (mg or µg) per 1000kcal, based on 110kcal/kg maintenance energy requirement (MER). FEDIAF guideline content for individual vitamins given as appropriate, for reference. Data are mean ± SD, analysed by Kruskal-Wallis NP ANOVA.
| **B-vitamin** | **FEDIAF**<br>(per 1000kcal) | **Meat-based**<br>(n=8) | **Plant-based**<br>(n=6) | **Renal**<br>(n=4) | **Statistics**<br>*P-value |
| --- | --- | --- | --- | --- | --- |
| Vitamin B1- Thiamine (mg) | 0.54 | 14.5 ± 22.6 | 3.58 ± 1.39 | 4.48 ± 2.67 | 0.89 |
| Vitamin B2- Riboflavin (mg) | 1.50 | 1.99 ± 0.97 | 1.30 ± 0.80 | 1.83 ± 0.91 | 0.33 |
| Vitamin B3- Niacin (mg) | 4.09 | 6.52 ± 3.43<sup>ab</sup> | 1.72 ± 0.81<sup>b</sup> | 25.4 ± 23.7<sup>a</sup> | 0.003 |
| Vitamin B5- Pantothenic acid (mg) | 3.55 | 5.79 ± 2.68 | 4.18 ± 2.27 | 35.1 ± 11.0 | 0.06 |
| Vitamin B6- Pyridoxine (mg) | 0.36 | 1.42 ± 1.27 | 0.73 ± 0.47 | 1.76 ± 0.43 | 0.07 |
| Vitamin B9- Folic acid (µg) | 64.5 | 111 ± 94<sup>ab</sup> | 41.3 ± 29.7<sup>b</sup> | 207 ± 32<sup>a</sup> | 0.02 |
| Vitamin B12- Cobalamin (µg) | 8.36 | 16.5 ± 9.6<sup>ab</sup> | 4.96 ± 4.54<sup>b</sup> | 25.5 ± 5.5<sup>a</sup> | 0.01 |
| Total B-vits (mg, 1-6) | - | 30.2 ± 26.8<sup>a</sup> | 11.5 ± 4.5<sup>b</sup> | 42.3 ± 28.0<sup>c</sup> | 0.05 |
| Total B-vits (µg, 9+12) | - | 127 ± 101<sup>a</sup> | 46.2 ± 32.4<sup>b</sup> | 233 ± 37<sup>c</sup> | 0.01 |

## Discussion

Adopting a plant-based dietary pattern is becoming increasingly common in Western society whether for health benefits (<u>Hall et al. 2021</u>; <u>Leonard et al. 2024</u>) or for environmental considerations (<u>Humpenöder et al. 2022</u>). Canines, as omnivores, are well-adapted to receive such a diet. Vegetarians and vegans, by definition “plant-based”, more commonly experience some micronutrient deficiencies (<u>Bath et al. 2024</u>; <u>Niklewicz et al. 2023</u>), easily rectified through supplementation. For companion animals fed a ‘nutritionally-complete’ plant-based diet, such deficiencies should not occur. The Food Standards Agency, UK requires labelling of foods as ‘complete’ to mean that feeding such food would give the companion animal all the nutrients it requires for maintenance or for growth and development. Few studies have independently tested this assumption. Here, we show that ‘complete’, dry plant-based foods for canines were replete in protein and amino acids, but consistently low in iodine and some B-vitamins – similar to observations made for human populations following a vegetarian or vegan diet (<u>Bath et al. 2024</u>; <u>Niklewicz et al. 2023</u>). Interestingly, guaranteed-analysis, veterinary-renal diets designed for a particular nutritional purpose – thus, low in protein to support dogs with moderate kidney disease – were also low in many essential amino acids.

### Protein, amino acids and fatty acids

Independent analysis of total crude protein correlated well with the sum of alpha amino acids measured in all foods, and the amount of protein as reported on food labels. The veterinary foods were, as expected, lower in measured total protein but, unexpectedly, were also relatively deficient (cf. guidelines for such foods) in a number of essential amino acids (EAA); 66% of the foods were low in at least one EAA despite us not reporting relatively low levels of methionine due to analytical considerations – that acid-hydrolysis can often result in an under-estimate of its concentration (<u>Gehrke et al. 1985</u>). Nevertheless, 9/10 of the EAAs for canines are reported: arginine, histidine, isoleucine, leucine, lysine, (methionine), phenylalanine, threonine and valine. One food had below the nutritional minimum guideline for 5 out of 9 EAAs tested. The longer-term effects of feeding such an EAA deficient diet to an animal with suspected CKD are not known. Current clinical guidelines for patients with CKD, without diabetes, advocate a diet low in protein but with supplemental keto-analogues of essential amino acids to support metabolic cycles dependent on supply of EAA (<u>Ikizler et al. 2020</u>). In canines, if the same diet were fed for a long-period, then it is possible that other co-morbidities might be exacerbated; low intake of S-containing amino acids, for example, is associated with an increased risk of developing dilated cardiomyopathy, due to reduced taurine synthesis (<u>Kaplan et al. 2018</u>).

It was hypothesised that plant-based foods would contain inadequate branched-chain amino acids (BCAAs; leucine, isoleucine and valine), as most dietary BCAAs are derived from meat, fish and dairy products (<u>Wolfe 2017</u>). However, all meat- and plant-based foods met minimum nutritional requirements for BCAAs and average concentrations were, in fact, greater in plant-based foods compared to those comprised of predominantly beef or lamb. Again, one of the veterinary-renal foods had lower than recommended BCAA content. Unlike cats, the majority of dogs can synthesise taurine endogenously using sulphur-containing amino acids (e.g. methionine and cysteine, (<u>Brosnan and Brosnan 2006</u>)). As such, taurine is not considered ‘essential’ for canines. For some large-breed dogs, such as Newfoundlands, taurine is essential, as a genetic mutation means they are unable to synthesis taurine endogenously and are therefore reliant on adequate dietary intake (<u>Li and Wu 2023</u>). Taurine might, therefore, be considered a conditionally-essential amino acid for some breeds of dog and dietary choices, such as breed-specific foods, for such breeds should be made on a case-by-case basis. Other factors such as nutrient-nutrient interactions, amount of dietary fibre and fat-to-protein ratio in the gastrointestinal tract may also affect the bioavailability of other marginal AAs, limiting their uptake, particularly in those foods with only marginally-replete content (<u>Mansilla et al. 2020</u>). It is almost churlish to propose, therefore, that for all foods designated as ‘complete,’ an assigned nutritional minimum for all essential and conditionally-essential amino acids should at least be followed (<u>Delaney and Fascetti 2012</u>).

### Fatty acids

Few nutritional recommendations exist for fatty acids (<u>FEDIAF 2022</u>). In this study, we were able to quantify n=36 fatty acids (from C6:0 to C22:6n3) and report that where guidelines exist, (on a g per total lipid mass basis) all foods were replete in fatty acids, including those essential for canines (e.g. C18:2n6, linoleic acid), despite the fact that the majority of n-3 long-chain fatty acids such as eicosapentaenoic acid (EPA, 20:5) and docosahexanenoic acid (DHA, 22:6) are traditionally sourced from marine oils (<u>Ahlstrøm et al. 2004</u>), which are not incorporated into plant-based foods. Alternative sources of omega 3- and 6 fatty acids for incorporation into plant-based foods, including chia and hemp seeds, flaxseed, walnuts, soya, seaweed, and microalgae, can be used satisfactorily (<u>TheVeganSociety 2023</u>). Hence, with the varied nutritional sources as listed in decreasing order of incorporation on the labels in the foods tested here, then all requirements for essential and non-essential fatty acids were adequately met.

### Major and trace minerals

Taken together, the current analysis of all major and trace elements in terms of compliance compared to relevant national guidelines for plant- and meat-based foods is similar to that reported previously (<u>Davies et al. 2017</u>). That is, foods were broadly compliant (n=342/391 [87%] of minerals were ‘in-range’), but only 16% of foods satisfied all mineral guidelines. Veterinary-renal foods where certain minerals are exempted (e.g., Ca and P) due to being low for a ‘particular nutritional purpose’ (‘PARNUTS’, (<u>Regulation 2020</u>)) were not included in this analysis and were balanced in terms of the Ca:P ratio. Plant-based foods, in general, contained greater potassium and magnesium, consistent with mineral enrichment within plants, as well as sufficient in elemental iron – which can, for example, be limiting for female vegetarians (<u>Pawlak, Berger, and Hines 2018</u>). In general, only sporadic deviations from recommended nutritional minimums or maximums were noted in some major minerals e.g. calcium, phosphorous and potassium and some trace (often selenium or zinc) minerals. It is unlikely therefore that any clinical signs of malnutrition would develop as a result of these micronutrient imbalances, unless fed exclusively for long periods of time and over multiple batches of the same food.

Nevertheless, of particular note was that iodine was below the guideline nutritional minimum in over half (57%, n=17 of 30) of all foods measured and remained when expressed on weight basis or corrected for energy density of foods. It was note-worthy that n=5 of 6 plant-based foods measured had relatively low iodine, which is in-keeping with people following plant-based diets (<u>Krajčovičová-Kudláčková et al. 2003</u>; <u>Bath et al. 2024</u>). It would be relatively simple to supplement these foods with plant-based sources of high iodine, such as seaweed or sea-kelp (<u>McCance and Widdowson 2014</u>). Indeed, the only plant-based food with adequate iodine had both seaweed and dried algae as significant ingredients. A previous study by us reported similar results for feline diets, which also varied considerably between batches for iodine content (<u>Alborough, Graham, and Gardner 2022</u>). Regardless, whilst iodine deficiency is common in vegetarians or vegans (<u>Eveleigh, Coneyworth, and Welham 2023</u>; <u>Weikert C and K. 2020</u>), few studies have reported any adverse effects of low iodine intake over the long-term in canines.

### Vitamin D

Vitamin D insufficiency has been previously reported in dogs fed commercial meat-based (<u>Sharp, Selting, and Ringold 2015</u>) or homemade diets (<u>Zafalon, Ruberti, et al. 2020</u>). Ingestion of food, as opposed to sunlight, is the primary source of Vitamin D for canines (<u>How, Hazewinkel, and Mol 1994</u>). In the current study, direct analysis of the foods for Vitamin D demonstrated none to be deficient. Vitamin D is a fat-soluble micronutrient with active endocrine properties (<u>Hurst, Homer, and Mellanby 2020</u>). The latter are particularly important during growth and development, given the important role Vitamin D has in calcium and phosphorus homeostasis; chronically low intake or deficiency of Vitamin D can cause bone de-mineralisation, through release of stored calcium and/or phosphate and may influence other non-skeletal related conditions (<u>Clarke, Hurst, and Mellanby 2021</u>).

Chronically elevated intake can lead to increased calcium and phosphorus absorption by the gut, with any subsequent hypercalcaemia being associated, in the longer-term, with chronic kidney disease (<u>Chacar et al. 2020</u>). Reporting values close to the upper nutritional guideline in pet food (one meat-based food was within 2% of the nutritional maximum) could become a problem if fed for a long period of time. Furthermore, considering the growth in plant-based food products for pets, it should also be noted that source of Vitamin D can influence bioavailability, which has not been measured here; active vitamin D2 (ergocalciferol) and D3 (1,25-dihydroxy-cholecalciferol) from meat-based sources are generally more bioavailable than vitamin-D2 from plant-based sources (<u>Tripkovic et al. 2012</u>; <u>Chungchunlam and Moughan 2023</u>).

### B-vitamins

B-vitamin deficiency can be caused by a number of gastrointestinal conditions in canines that, irrespective of dietary sufficiency, mean that B-vitamin uptake in the gut is reduced and less are bioavailable for cellular functions (<u>Kather et al. 2020</u>). B-vitamin deficiency can have a range of effects on the body, including but not limited to, affecting the integrity of the dermal layer, acute disturbances of the central nervous system, lethargy, vomiting and diarrhoea (<u>Gildea et al. 1935</u>; <u>Kather et al. 2020</u>). Research into B-vitamin homeostasis in companion animals is lacking and complicated by variation according to breed (<u>Kather et al. 2020</u>). B-vitamins are water-soluble and readily excreted in urine if taken in excess – which is common if predominantly consuming an animal-based diet. Hypervitaminosis, particularly of B-vitamins, is therefore rarely of clinical concern. In contrast, deficiency is commonly reported in vegetarians and vegans (<u>Niklewicz et al. 2023</u>). Regardless, all B-vitamins can easily be obtained from plant-based sources, although may need to be consumed in higher quantities due to poor bioavailability (<u>Chungchunlam and Moughan 2023</u>). Consequently, humans often rely, at least partially, on supplementation from fortified foods, which may or may not be sufficient to meet dietary requirements (<u>Niklewicz et al. 2023</u>). A recent study found that Vitamin B12 status was similar between vegans (almost all of whom consumed supplements) and non-vegans (approx. 1/3 consumed supplements) (<u>Weikert et al. 2020</u>). For most foods analysed in this study – particularly plant-based, where the majority were lower in B1, B2, B3, B5, B9 and B12 – then further supplementation is recommended. Indeed, even for meat-based foods, B-vitamin supplementation using pre-mixes is common due to variability in B-vitamin content between animal products (e.g. muscle, organs) which can vary; cobalamin (vitamin B12) is low in muscle tissue for example (<u>Pinchen et al. 2020</u>). In addition, possible losses of B-vitamins (e.g. A, D, E, C and B9 [folic acid]) can occur during the refinement process toward production of a dog kibble (<u>Morin, Gorman, and Lambrakis 2021</u>). Increased temperature, for example, reduces active B-vitamin content in extruded foods (<u>Morin, Gorman, and Lambrakis 2021</u>).

### Overall compliance of pet foods

Finally, all 31 complete dry dog foods were tested singularly and compared to European Pet Food Industry Federation (FEDIAF) guidelines (FEDIAF, 2021). When compared to FEDIAF guidelines, 17/31 foods tested met all amino acid, 5/31 met all mineral, 4/18 met B-vitamin and all tested met vitamin D guidelines. No food met FEDIAF guidelines for all nutrients. It is important to note that this analysis was conducted on complete, pre-digested food. There are many factors that will influence the uptake of nutrients within the body. Nevertheless, even nutrients that are replete in food may become deficient or have low bioavailability/bio-accessibility in the gastrointestinal tract due to nutrient-nutrient interactions.

### Conclusion and future directions

Plant-based feeding of companion canines is becoming increasingly frequent. It is important to reassure owners/guardians of such pets that the feeding of a ’complete’ food, as designated on the label, is replete in all nutrients and micronutrients that are considered essential for dogs. Our study reports the first complete nutritional comparison of meat-based (including veterinary diets) and plant-based foods for canines in the UK and suggests variable compliance: 55%, 100%, 16%, 100% and 24% of foods met nutritional guidelines for amino acid, fatty acids, major and trace mineral, vitamin D and B-vitamins, respectively. Plant-based foods were often low in iodine and many of the B-vitamins, which could be easily corrected by incorporation of mineral-rich ingredients and/or supplementation. Veterinary diets with lower protein content by design (e.g. for dogs with kidney disease) often had below-recommended levels of incorporation of essential amino acids, which could also be easily corrected through supplementation with keto- analogues of essential amino acids. This study has only compared foods for adult dogs. Arguably, such micronutrient deficiencies would have greater impact if fed during growth & development or reproductive phases, when greater demand is placed on metabolic partitioning of nutrients. Further analyses of such foods, where available, is warranted. Many foods were compliant, but marginal, with respect to nutrient composition. In this instance, further factors such as bioaccessibility, digestibility and nutrient-nutrient interactions may produce systemic micronutrient deficiencies, despite guidelines taking these factors into account when establishing guidelines (<u>Officials) 2014</u>; <u>FEDIAF 2022</u>). Clearly, analysis of such multi-variate, gastrointestinal interactions is beyond the scope of the current study, but when novel foods are brought to market that might evidently have bioaccessibility effects (e.g. plant-based foods), then such studies are warranted.

## Author contributions

R.A.B and D.S.G designed the research; R.A.B, D.L and R.B conducted the research; R.A.B and D.S.G analysed the data; D.S.G gained the funding and supervised the project; R.A.B and D.S.G co-wrote the manuscript. Both authors critically evaluated the paper and take responsibility for its final content.

## Acknowledgements and Financial Disclosure

The authors gratefully acknowledge the help of Dr Catherine Williams (NUVetNA Ltd) and Saul Velasquez for assistance with mineral analyses, Noriane Cochetal for assistance with amino acid data analysis, Drs Louise Williams and Jon Stubberfield for macronutrient analysis. This study was funded by the Biotechnology and Biological Sciences Research Council (BBSRC) as part of the University of Nottingham DTP PhD studentship awarded to R.A.B (Grant code: RS86P5).

DSG conducts nutritional analysis for one of the tested companies’ foods. D.S.G is not involved in the formulation of the food, nor has influence on the design or reporting of results. Since completing the analysis, R.A.B has purchased shares in the same company; testing was completed before this occurred.

