## Supplementary Information for "Nutritional analysis of commercially available, complete plant- and meat-based dry dog foods in the UK"

**Table S1**. Standard Reference Material (SRM) transitions used for the AA standards, calibration standards and amino acid hydrolysate samples.

**Table S2.** uHPLC-MS/MS Instrumentation, Chromatographic and MS Parameters uHPLC Gradient for analysis of individual amino acids

**Table S3**. Serine (g/100g DM) as determined in 30 of 31 samples of pet feed

**Table S4.** Preparation of buffers for extraction of cytoplasmic (i.e. free) lipid from feeds using the ‘sucrose cushion’ method

**Table S5**. LC-MS methods & conditions for determination of Vitamin D2 and D3**.**

**Figure S1.** Example trace for detection of Vitamin B12 in extracted pet food samples

**Table S6.** LoQ values for B vitamin analysis

**Table S1**. Standard Reference Material (SRM) transitions used for the AA standards, calibration standards and amino acid hydrolysate samples.

| Compound | Precursor (m/z) | Product (m/z) | Collision Energy (V) | Compound | Precursor (m/z) | Product (m/z) | Collision Energy (V) |
|---|---|---|---|---|---|---|---|
| Alanine | 90.1 | 44.1 | 10 | Alanine 13C15N | 94.1 | 47.1 | 10 |
| Alanine | 90.1 | 90.1 | 5 | Alanine 13C15N | 94.1 | 94.1 | 5 |
| Arginine | 175.1 | 70.1 | 22 | Arginine 13C15N | 185.2 | 75.1 | 22 |
| Arginine | 175.1 | 116 | 14 | Arginine 13C15N | 185.2 | 125.1 | 14 |
| Asparatate | 134 | 74.1 | 15 | Asparagine 13C15N | 139.1 | 77.1 | 10 |
| Asparatate | 134 | 88.1 | 10 | Asparagine 13C15N | 139.1 | 59.1 | 10 |
| Cysteic acid | 170.1 | 129 | 4 | Aspartate 13C15N | 139.1 | 92.2 | 10 |
| Cysteic acid | 170.1 | 170.1 | 4 | Aspartate 13C15N | 139.1 | 77.1 | 15 |
| Cystine | 241 | 74.2 | 27 | Cysteine 13C15N | 126.1 | 79.1 | 10 |
| Cystine | 241 | 152 | 13 | Cysteine 13C15N | 126.1 | 45.1 | 10 |
| Glutamic acid | 148.1 | 84.1 | 15 | Glutamate 13C15N | 154.1 | 89.1 | 15 |
| Glutamic acid | 148.1 | 129.9 | 10 | Glutamate 13C15N | 154.1 | 89.1 | 10 |
| Glycine | 76 | 30.2 | 10 | Glutamine 13C15N | 154.1 | 74.1 | 10 |
| Glycine | 76 | 76.1 | 5 | Glutamine 13C15N | 154.1 | 95 | 10 |
| Histidine | 156.1 | 83.1 | 24 | Glycine 13C15N | 79.1 | 79.1 | 10 |
| Histidine | 156.1 | 110.1 | 13 | Glycine 13C15N | 79.1 | 61.1 | 5 |
| Isoleucine | 132.1 | 69.1 | 16 | Histidine 13C15N | 165.1 | 118.1 | 24 |
| Isoleucine | 132.1 | 86.1 | 10 | Histidine 13C15N | 165.1 | 91.5 | 13 |
| Leucine | 132.1 | 43.2 | 24 | Isoleucine 13C15N | 139.1 | 74.1 | 16 |
| Leucine | 132.1 | 44.1 | 22 | Isoleucine 13C15N | 139.1 | 92.2 | 10 |
| Leucine | 132.1 | 86.1 | 10 | Leucine 13C15N | 139.1 | 46.1 | 24 |
| Lysine | 147.1 | 84.1 | 16 | Leucine 13C15N | 139.1 | 47.1 | 22 |
| Lysine | 147.1 | 130.1 | 10 | Leucine 13C15N | 139.1 | 92.1 | 10 |
| Methionine | 150.1 | 104.1 | 10 | Lysine 13C15N | 155.1 | 90.2 | 16 |
| Methionine | 150.1 | 133 | 10 | Lysine 13C15N | 155.1 | 137.2 | 10 |
| Methionine sulfone | 182.2 | 56.1 | 15 | Methionine 13C15N | 156.1 | 109 | 10 |
| Methionine sulfone | 182.2 | 136.1 | 15 | Methionine 13C15N | 156.1 | 139.1 | 10 |
| Phenylalanine | 166.1 | 103.1 | 27 | Phenylalanine 13C15N | 176.2 | 107.1 | 27 |
| Phenylalanine | 166.1 | 120.1 | 11 | Phenylalanine 13C15N | 176.2 | 129.1 | 11 |
| Proline | 116.1 | 43.2 | 28 | Proline 13C15N | 122.1 | 46.3 | 28 |
| Proline | 116.1 | 70 | 15 | Proline 13C15N | 122.1 | 75.1 | 15 |
| Serine | 106.1 | 60.3 | 10 | Serine 13C15N | 110.1 | 63 | 10 |
| Serine | 106.1 | 88 | 10 | Serine 13C15N | 110.1 | 83.1 | 10 |
| Threonine | 120.1 | 74.1 | 10 | Threonine 13C15N | 125.1 | 78.1 | 10 |
| Threonine | 120.1 | 102 | 10 | Threonine 13C15N | 125.1 | 107.1 | 10 |
| Tyrosine | 182.1 | 136.1 | 12 | Tryptophan 13C15N | 218.3 | 156.2 | 10 |
| Tyrosine | 182.1 | 165 | 10 | Tryptophan 13C15N | 218.3 | 46.1 | 10 |
| Valine | 118.1 | 55.1 | 19 | Tryptophan 13C15N | 218.3 | 70.1 | 10 |
| Valine | 118.1 | 72.1 | 10 | Tyrosine 13C15N | 192.2 | 145.2 | 12 |
| | | | | Tyrosine 13C15N | 192.2 | 174.2 | 10 |
| | | | | Valine 13C15N | 124.1 | 59 | 19 |
| | | | | Valine 13C15N | 124.1 | 77.1 | 10 |

**Table S2.** uHPLC-MS/MS Instrumentation, Chromatographic and MS Parameters uHPLC Gradient for analysis of individual amino acids

| Time (min) | Flow Rate (ml/min) | %A | %B |
|---|---|---|---|
| 0 | 0.3 | 100 | 0 |
| 5 | 0.3 | 100 | 0 |
| 7 | 0.3 | 0 | 100 |
| 14 | 0.3 | 0 | 100 |
| 14.5 | 0.35 | 100 | 0 |
| 16.5 | 0.35 | 100 | 0 |
| 17 | 0.3 | 100 | 0 |
| 18 | 0.3 | 100 | 0 |

**Table S3**. Serine (g/100g DM) as determined in 30 of 31 samples of pet feed

| SampleID | Meat-based | Plant-based | Renal |
|---|---|---|---|
| S#265 | 8.60 | | |
| S#266 | 10.85 | | |
| S#267 | 11.40 | | |
| S#268 | 14.71 | | |
| S#269 | 11.82 | | |
| S#270 | 12.47 | | |
| S#271 | 8.31 | | |
| S#272 | 14.83 | | |
| S#273 | 6.73 | | |
| S#274 | 11.78 | | |
| S#275 | 5.80 | | |
| S#276 | 12.01 | | |
| S#277 | 10.76 | | |
| S#278 | 10.23 | | |
| S#286 | 10.57 | | |
| S#291 | 10.59 | | |
| S#292 | 11.81 | | |
| S#293 | 11.33 | | |
| S#294 | 9.94 | | |
| S#288 | | 12.60 | |
| S#289 | | 11.97 | |
| S#290 | | 9.18 | |
| S#284 | | 12.32 | |
| S#285 | | 6.55 | |
| S#259 | | 6.86 | |
| S#279 | | | 7.30 |
| S#280 | | | 5.88 |
| S#281 | | | 26.96 |
| S#282 | | | 6.32 |
| S#283 | | | 7.70 |
| **Mean (SD)** | **10.7 (2.3)** | **9.9 (2.7)** | **10.8 (9.0)** |

**Table S4.** Preparation of buffers for extraction of cytoplasmic (i.e. free) lipid from feeds using ‘sucrose cushion’ method

a) Composition of Isolation buffer:

| Ingredient | Lot | Dry weight (g) |
|---|---|---|
| 50mM HEPES | 7M015925 | 11.9266 |
| 10mM Potassium chloride | SLBX747 | 0.7463 |
| 62.5mM Potassium acetate | 010409BH | 6.1472 |
| 5mM EGTA | 059K54251 | 1.915 |
| 5mM Dithiothreitol (DTT) | 00638353 | 0.7738 |
| 1mM Magnesium chloride | 2407C173 | 0.0956 |
| 5mM EDTA | 176442 | 1.8639 |

Make up to 1L using deionized water. pH adjusted to 7.5 using 10M NaOH.

b) 0.6M Sucrose extraction buffer:

| Ingredient | Lot | Dry weight (g) |
|---|---|---|
| Protease-free sucrose | 18A2756188 | 20.5025 |
| 1% w/v Bovine serum albumin (BSA) | 1703030465 | 0.999 |

Make up to 100mL with isolation buffer (S3a)

c) 0.25M Sucrose extraction buffer:

| Ingredient | Lot | Dry weight (g) |
|---|---|---|
| Protease-free sucrose | 18A2756188 | 8.4957 |
| 1% w/v Bovine serum albumin (BSA) | 1703030465 | 1.0045 |

Make up to 100mL with isolation buffer (S3a)

**Table S5**. **LC-MS methods & conditions for determination of Vitamin D2 and D3.**

The MRM transitions are listed below.

| Compound | Precursor m/z | Product m/z | Dwell (s) | Cone (V) | Collision (V) |
|---|---|---|---|---|---|
| Vitamin D3 Quant | 560.3 | 298.1 | 0.02 | 43 | 19 |
| Vitamin D3 Qual | 560.3 | 161.0 | 0.02 | 43 | 36 |
| d3-Vitamin D3 Quant | 563.3 | 301.2 | 0.02 | 43 | 19 |
| Vitamin D2 Quant | 572.3 | 298.1 | 0.02 | 43 | 19 |
| Vitamin D2 Qual | 572.3 | 311.8 | 0.02 | 43 | 15 |
| d3-Vitamin D2 Quant | 575.3 | 301.2 | 0.02 | 43 | 19 |

4.5 ml of 1.5 M potassium hydroxide in ethanol containing 0.5% w/v pyrogallol was added to all samples, which were then vortexed, heated to 80°C for 1 hour, with periodic mixing. Once cool, 3 ml of a 1% w/v potassium chloride solution was added, followed by 3 ml of hexane. Samples were then vortexed, followed by centrifugation at 1400 rpm for 5 minutes to sediment particulate matter. The upper hexane layer was removed into clean glass tubes. A further hexane extraction was carried out on the saponified material, which was then pooled with the first hexane extraction. The hexane extracts were then dried under nitrogen and resuspended in 500 µl of ethyl acetate, with samples transferred to 1.5 ml Eppendorf tubes. The ethyl acetate was evaporated under nitrogen. 120 µl of a 1 mg/ml PTAD (4-phenyl-1,2,4-triazoline-3,5-dione) solution in acetonitrile was added to samples, which were left to react for 1 hour in the dark at room temperature, followed by the addition of 40 µl of water. Samples were vortexed then centrifuged at 10,000 rpm for 5 minutes and the supernatant transferred to low volume, pulled point glass vials for LC-MS analysis. Samples were analysed on a Waters Xevo TQ-S mass spectrometer coupled to a Waters Acquity I class UPLC with an Acquity UPLC BEH C18 column (2.1 x 100 mm, 1.7 µm) and a 5 mm guard column attached. The UPLC mobile phases consisted of; solvent A, 90:10 water/acetonitrile with 0.1% v/v formic acid and solvent B, 100% methanol with 0.1% v/v formic acid. The column temperature was maintained at 50 °C, with the flow rate set at 0.4 ml/min. The column elution gradient had an initial composition comprising 50% A and 50% B. This was held for 1 min, then ramped up gradually to 72% B over a period of 7 mins. A further ramp to 100% soln B was completed in 2 mins, held for a further 2 mins. The gradient was then ramped back to 50% B over 3 mins and equilibrated for a further 2 minutes at the starting conditions. The mass spectrometer source settings were as follows: A capillary voltage of 3 kV in positive mode and a source offset of 30 V. The desolvation temperature was set to 300 °C, with a desolvation flow rate of 800 L/Hr and cone flow rate of 150 L/Hr.

**Figure S1.** Example trace for detection of Vitamin B12 in extracted pet food samples

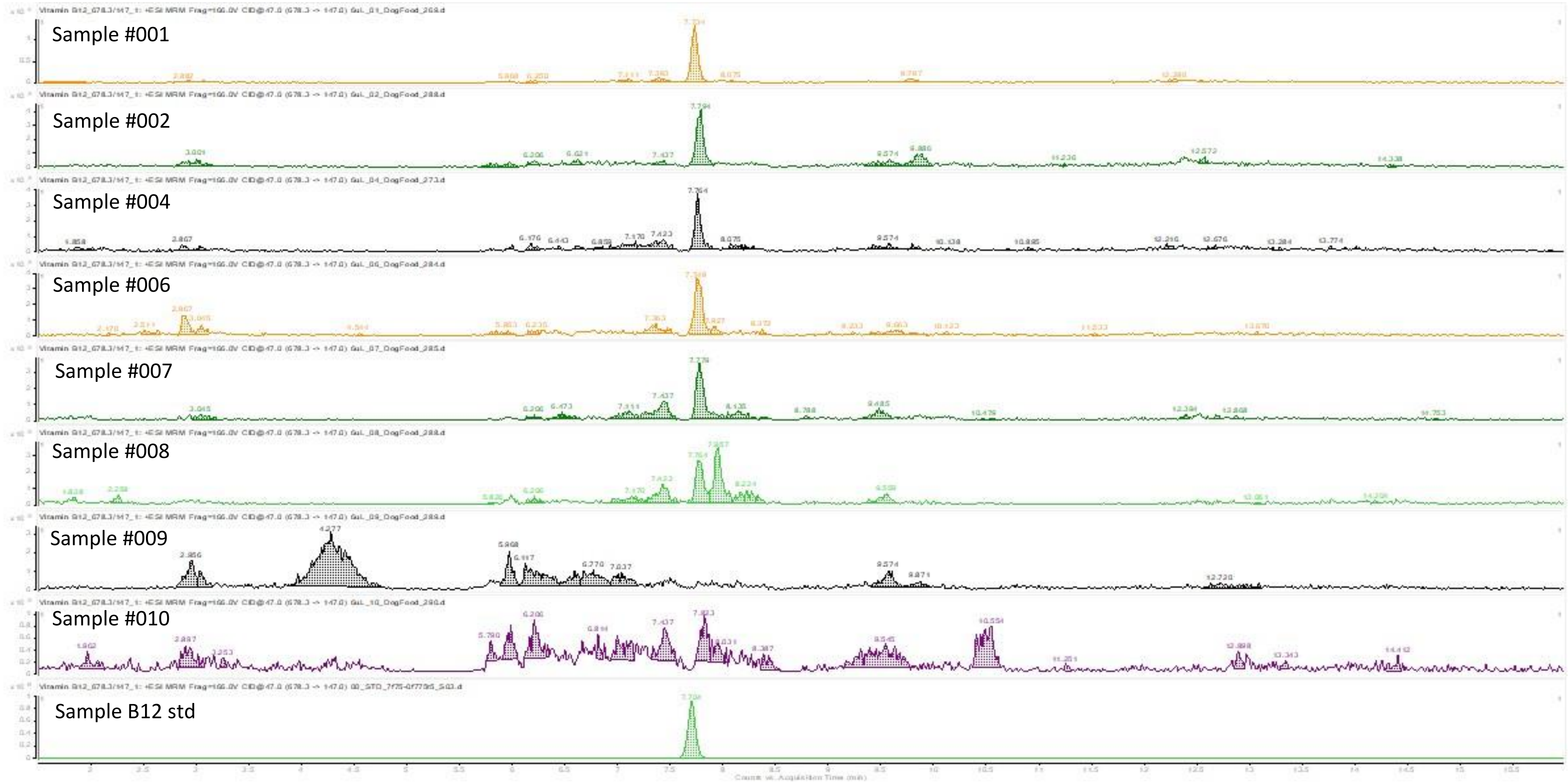


Detection of B12 in dog food samples. Samples #001 - #008, B12 detectable. Samples #009, #010 B12 not reliably detectable.

**Table S6.** LoQ values for B vitamin analysis

| B-vitamin | LoQ (mg/1000kcal) |
|---|---|
| Vitamin B1- Thiamine | 0.00159213 |
| Vitamin B2- Riboflavin | 0.00056454 |
| Vitamin B3- Nicotinamide | 0.00018318 |
| Vitamin B3- Niacin | 0.000061555 |
| Vitamin B5- Pantothenic acid | 0.000109615 |
| Vitamin B6- Pyridoxal | 0.00008358 |
| Vitamin B6- Pyridoxamine | 0.000252294 |
| Vitamin B6- Pyridoxine | 0.00008459 |
| Vitamin B7- Biotin | 0.000366465 |
| Vitamin B9- Folic acid | 0.11035 |
| Vitamin B12- Cobalamin | 0.203307 |
